# Reversal learning performance in the XY* mouse model of Klinefelter and Turner Syndromes

**DOI:** 10.1101/672022

**Authors:** Shawn M. Aarde, Haley Hrncir, Arthur P. Arnold, J. David Jentsch

## Abstract

Klinefelter syndrome (KS; 47, XXY) and Turner syndrome (TS; 45, XO) are caused by two relatively common sex chromosome aneuploidies. These conditions are associated with an increased odds of neuropsychiatric disorders, including attention deficit/hyperactivity disorder (ADHD), as well as impairments in cognition that include learning delays, attentional dysfunction and impulsivity. We studied cognitive functions in the XY* mouse model, which allows comparison of XXY to XY males (KS model), and XO to XX females (TS model). We evaluated adult mice with and without gonads, using a version of an operant reversal-learning task (RLT) that can be used to measure various facets of learning, impulsivity and attention. In the KS model, only one measure related to impulsivity – perseverative responding under reversal conditions – reliably discriminated gonadally intact XXY and XY mice. In contrast, a fundamental learning impairment (more trials to criterion in acquisition phase) in XXY mice, as compared to XY, was observed in gonadectomized subjects. No other task measures showed differences consistent with KS. In the TS mouse model, XO mice did not show a pattern of results consistent with TS, similar to past observations. Thus, the application of this RLT to these XY* models reveals only limited behavioral impairments relevant to KS.

## 1 Introduction

Sex-chromosome aneuploidies, such as Klinefelter syndrome (KS; 47, XXY) and Turner syndrome (TS; 45, X), are associated with various cognitive impairments and with an increased risk of psychiatric and/or neurodevelopmental disorders, including attention deficit-hyperactivity disorder, autism, schizophrenia, and bipolar disorder (Belling et al., 2017; Cederlöf et al., 2014; Russell et al., 2006; Tartaglia et al., 2017; Zhao and Gong, 2017). For example, an increased risk of ADHD has been observed in people with either KS (Cederlöf et al., 2014) or TS (Russell et al., 2006). Moreover, KS men who do not have a diagnosis of autism spectrum disorder nonetheless have higher Autism-spectrum Quotient scores (AQ) (Baron-Cohen et al., 2001), including poorer scores on the “attention switching” subscale, indicating subclinical neurobehavioral manifestations that will require quantitative, rather than diagnostic, behavioral approaches (Cederlöf et al., 2014; van Rijn et al., 2012b, 2014).

Cognitive assessments indicate a modestly lower full-scale IQ (5-10 points) in both KS and TS, with higher performance IQ relative to verbal IQ in KS, and the opposite relationship in TS (Hong and Reiss, 2014). Moreover, in a recent assessment that included a Stroop Word-Color Test and the Wisconsin Card Sorting Test, KS men exhibited an greater Stroop effect (thus, poorer inhibition) and made more errors (including more perseverative errors) on the card sort task relative to controls (thus, poorer behavioral flexibility) (Skakkebæk et al., 2014).

To better understand the mechanisms by which sex-chromosome aneuploidies impact phenotypes, various animal models of KS or TS have been developed (Burgoyne and Arnold, 2016; Cox et al., 2014; Lue et al., 2010; Wistuba, 2010). Behavioral studies of these models indicate that both KS and TS aneuploidies impact learning, impulsivity and/or attention (Chen et al., 2013b; Cox et al., 2015; Davies et al., 2005, 2007; Isles et al., 2004; Lewejohann et al., 2009; Lue et al., 2005).

The particular KS and TS models that we used were derived from the XY* mouse. In this mouse, the Y chromosome has a modified pseudoautosomal region that recombines abnormally with the X chromosome. This generates four male gametes: X, Y*, X^Y*^ (fused X and Y*), and Y^*X^ (which paradoxically is not a Y chromosome, but predominantly the pseudoautosomal region of the X chromosome) (Burgoyne and Arnold, 2016). Thus, mating an XY* male to an XX female produces mice that are the near-equivalent of XX, XY (i.e., XY*), XXY (i.e., 40,XX^Y*^), and XO (i.e., 40,XY^*X^) (Burgoyne et al., 1998; Burgoyne and Arnold, 2016; Eicher et al., 1991; Wistuba et al., 2010). Although 39,XO mice have been generated and studied (Davies et al., 2005, 2007; Isles et al., 2004; Lopes et al., 2010; Raznahan et al., 2013), the current study used 40,XO mice (i.e., 40,XY^*X^) on a C57BL/6J background, because the 39,XO model is not available on that background.

Although the data are limited, mouse models of KS do appear to have some face validity in terms of learning impairments (Ross et al., 2008; Rovet et al., 1996; Skakkebæk et al., 2017). For example, in a novel-object recognition test, XXY mice from the XY* model failed to show the typical pattern of more exploration of the novel object relative to the more familiar object, but control XY mice did (Lewejohann et al., 2009). In a different KS model, 41,XXY mice were slower to acquire Pavlovian appetitive approach behavior than XY mice (Lue et al., 2005).

Regarding TS mouse models, performance on a serial reversal-learning task (Y-maze-based, visual, nonspatial) was worse in 39,XO than 40,XX mice, but only when the X-chromosome in the 39,XO mice was of maternal origin (Davies et al., 2005). Thus, 39,XO mice possessing the maternal X have face validity for the general learning impairments observed in TS (Garron, 1977; Loesch et al., 2005; Mazzocco, 2006). However, as the relevance of imprinting to learning impairments in TS remains unclear (Lepage et al., 2012), so does its relevance to the validity of mouse models of TS (Davies et al., 2005, 2007).

On the 5-choice serial reaction time task (5-CSRTT; tests visual attention), 39,XO mice displayed poorer accuracy and slower reaction times compared to 40,XX mice (outbred MF1 strain background) (Davies et al., 2007). However, the presence of the Y*^X^ chromosome in 40,XO mice rescued the attention deficit and there were no parent-of-origin effects detected. Additionally, on measures of impulsivity, 39,XO and 40,XX mice did not differ. Thus, the 39,XO genotype appears to have face validity in terms of the attentional deficits observed in TS (Delooz et al., 1993; Green et al., 2018; Mauger et al., 2018; McCauley et al., 1987; Ross et al., 2002; Russell et al., 2006), but not for the impulsivity observed in TS (Romans et al., 1997; Russell et al., 2006; Tamm et al., 2003).

To further study behavior and cognition in the XY* model, we compared XXY males to XY males (KS comparison), as well as XO females (40,XO; i.e., XY*^x^) to XX females (TS comparison), on an operant reversal-learning task (RLT) (Laughlin et al., 2011; Linden et al., 2018). This task was chosen because it measures some of the constructs impaired in KS and TS (learning, impulsivity, and attention), as well as for its relevance to the current literature.

A recent advance in the behavioral analysis of this particular version of the reversal learning task (Laughlin et al., 2011; Linden et al., 2018) makes use of *a priori* defined and empirically determined “change points” in the learning curve to better resolve within-phase behavior dynamics (Gallistel et al., 2004; Linden et al., 2018; Papachristos and Gallistel, 2006). In feedback-based operant-learning tasks, the acquisition curve of an individual subject is typically not one of a negatively accelerating, gradual progress toward a performance asymptote (Gallistel et al., 2004; Papachristos and Gallistel, 2006). Rather, such learning curves are typically an artifact of group averaging and thus fail to represent the behavioral dynamics of individuals which include changes in the rate of learning caused by prolonged intervals of stable performance and/or abrupt, large step-like changes in proficiency – i.e., “change points” (Gallistel et al., 2004; Papachristos and Gallistel, 2006).

However, a recent report made successful use of these change points to analyze data from a spatial reversal-learning task in rats (Klanker et al., 2015). Specifically, they split trials into those occurring before (PRE) or after (POST) the largest empirically-determined change point in the individual learning curves. They observed that, for rats that learned the reversal, phasic post-reward dopamine release in the ventromedial striatum was lower POST than PRE change-point. Additionally, post-cue dopamine release was higher on trials that followed a rewarded trial than on those prior rewarded trials – but, only PRE change-point. Thus, we made use of this analytical tool as it appears to better resolve discrete phases in learning.

Some effects of KS and TS are potentially attributed to altered levels of gonadal hormones. KS men have lower levels of testosterone than XY men (Gravholt et al., 2018; Klein et al., 2018), and TS women have altered ovarian function (Klein et al., 2018; Romans et al., 1998; Ross et al., 1995, 2004; Rovet, 2004). Accordingly, we analyzed the role of gonadal hormones on task performance by comparing gonadectomized mice to gonad-intact controls.

Under the assumptions that the XXY and XO genotypes of the XY* mice are sufficiently valid models of the genotypes of KS and TS, respectively, we predicted a pattern of genotype effects on behavior similar to that seen in people with KS and TS, with the exception of premature responding in the XO vs XX comparison for which no difference was predicted based upon prior performance on the 5-CSRTT/1-CSRTT (Davies et al., 2007). We expected that if genotypes differed on behavioral measures, that some of the differences might disappear if they depended on group differences in levels of gonadal hormones in any life stage. However, sex-hormone replacement therapies in adulthood do not appear to remediate the cognitive deficits observed in people with KS (Fales et al., 2003; Kompus et al., 2011; Liberato et al., 2017; Skakkebæk et al., 2017) or TS (Klein et al., 2018; Ross et al., 2002, 2004), and the cognitive consequences of reducing/blocking sex hormones in people with either KS or TS, or in this animal model, are unknown.

## 2 METHODS

### 2.1 Subjects

#### 2.1.1 XY* (XY-star) mouse model

Subjects were adult (100-129 days old at the time of testing) mice from the XY* model (Burgoyne et al., 1998; Burgoyne and Arnold, 2016; Eicher et al., 1991), backcrossed from Jackson lab strain 2021 to C57BL/6J for at least 13 generations. Offspring were produced by mating XY* males to XX females. This model produces 4 genotypes – XY*^X^, XX, XY* and XXY* – that are the near-equivalent of XO, XX, XY, and XXY, respectively (Burgoyne and Arnold, 2016; Chen et al., 2013a; Cox et al., 2014). The Y*^X^ chromosome is an X chromosome with massive deletion of most genes, leaving the pseudoautosomal region and a few nearby genes (Burgoyne and Arnold, 2016), so that XY*^X^ females have X monosomy (40, XO) except for the presence of a second PAR. In this model, XY and XXY mice have testes, and XO and XX mice have ovaries. In XXY (40, XXY) mice, one X chromosome and the Y chromosome are fused end-to-end (Burgoyne and Arnold, 2016).

Gonadectomy, or a control sham surgery, were performed at 72-99 days of age (mean=82), and behavioral testing began 25-41 (mean=30) days later. The 8 groups were: XY,GDX (N = 12); XY,SHAM (N = 11); XXY,GDX (N = 15); XXY,SHAM (N = 16); XX,GDX (N = 13); XX,SHAM (N = 14); XO,GDX (N = 14); XO,SHAM (N = 14).

#### 2.1.2 Genotyping

Genotype was determined by combined measurements of anogenital distance and fluorescent *in-situ* hybridization to detect X and Y chromosomes in interphase lymphocytes using the Kreatech KI-30505 kit (Leica Biosystems). Males also had their genotype verified by visual assessment of testis size; males with one X chromosome have testes roughly 6 times larger than males with two X chromosomes (Wistuba, 2010).

#### 2.1.3 Husbandry

After gonadectomy, mice were transferred to a vivarium that varied in temperature (69-79°F) and humidity (∼20-∼70%), under a 14h:10h light:dark cycle. Behavioral testing was performed 1-4 h before onset of the dark phase. Mice were housed in groups of 2-4 mice/cage, on sawdust bedding.

To facilitate motivation to perform the task, all mice were food-restricted to maintain a body weight of 80-85% of baseline (pre-restriction) weights. The daily allocation of standard rodent chow was fed about 30 min after testing, adjusted daily based on overnight changes in body weight, the difference between current and target weight (82.5%), and the amount of food consumed during testing. Water was always available except during testing (∼1 h), and the post-testing interval before being given their chow. All experimental procedures were approved by the UCLA Chancellor’s Animal Research Committee.

### 2.2 Operant Conditioning Testing - Reversal-Learning Task (RLT)

#### 2.2.1 Overview, Equipment

Operant chambers (Med Associates; St. Albans VT) were housed within sound-attenuating cubicles and were equipped with: 5 horizontally arranged nose-poke apertures on one curved wall, a reinforcer-delivery magazine on the opposite wall, a white-noise generator (∼85 dB; always on), and a house light (located outside and above the magazine-side of the chamber). Nose-poke apertures and the magazine were illuminated via recessed lamps, and entries into apertures/magazine were detected via interruption of an infrared-beam sensor. Data were collected via MedPC IV software (Med Associates) running custom behavioral programs.

#### 2.2.2 Habituation Phase

After initiation of food restriction, mice were transferred to the operant testing room where the equipment was turned on. Mice were placed singly into clean home cages (no bedding, two nestlets). After ∼ 30 min, a ceramic ramekin containing 1.4 g of the reward (14-mg Dustless Precision purified reinforcer pellets; BioServ item #F05684) was placed into the cage. After 1 h, the mice were returned to their original home cages. This was repeated for a total of 4 consecutive days.

#### 2.2.3 Magazine Training

Mice were next given 2 sessions of magazine training, during which reinforcer pellets were delivered on a modified variable-time schedule (30 ± 0, 5, or 10 s) in which reinforcers were delivered every ∼30 s, so long as a nose-poke into the magazine was detected before the time of the next pellet delivery (i.e., cessation of magazine poking suspended the timer and thus stopped further pellet deliveries until poking resumed). The magazine was illuminated when the pellet was delivered and then darkened when an entry into the magazine was detected. Sessions ended after delivery of 60 pellets or after 60 min, whichever occurred first.

#### 2.2.4 Observing-Response Training

Mice were next trained to produce a sustained duration nose-poke response into the central aperture (hole 3 of 5 within the horizontal array); this was termed an “observing” response, for use in the acquisition and reversal phases of testing. If the mice were successful at sustaining their response for the required duration, a pellet was delivered into the magazine, the magazine light was illuminated, and the center aperture was darkened. Failure to sustain the response caused a 2-s timeout during which the house light was illuminated and the center aperture was darkened. The required holding duration (RHD) was block randomized and possible durations were incremented in the next session if the mouse met performance criterion as follows (failure to meet criterion resulted in a repeat of the condition):

a. RHD = 1, 5, or 10 csec, criterion >= 30 pellets in < 60 min
b. RHD = 10, 20, or 30 csec, criterion >= 40 pellets in < 60 min
c. RHD = 20, 30, or 40 csec, criterion >= 40 pellets in < 60 min
d. RHD = 30, 40, or 50 csec, criterion >= 50 pellets in < 60 min

#### 2.2.5 Acquisition Learning Phase (ACQ)

Mice were next trained in ∼1-h daily sessions to acquire a simple spatial discrimination. Each trial began with illumination of hole 3 of 5; upon completion of the observing response (RHD = 1, 10 or 20 csec), the center aperture was darkened and the flanking apertures (holes 2 and 4 of 5) were illuminated. For each mouse, one of the two holes was *a priori* assigned as the correct side (designated correct aperture was counterbalanced across groups), and responses into that aperture would lead to pellet delivery into an illuminated magazine. Reward retrieval would darken the magazine and trigger the start of a 3-s inter-trial interval (ITI) after which the center aperture was re-illuminated and the next trial commenced.

Responses into the designated incorrect aperture triggered a 5-s timeout, during which all aperture lights were extinguished and the house light was illuminated. Alternatively, failure to nose-poke into either of the lit target apertures within 30-s of their illumination was counted as a response omission and led to a 5-sec timeout. Timeouts for incorrect responses and omissions were followed by a 3-sec ITI. Timeouts for observing-response failures continued as scheduled in observing-response training. However, after these timeouts, the ACQ trial continued until a successful observing-response triggered target presentation.

Premature responses (responses to the target apertures *before* target presentation) and extraneous observing responses (EOR – repetitive responses into the central aperture *during* target presentation) were recorded but had no scheduled consequences.

Subjects continued daily testing until they reached a performance criterion of 80% correct within a sliding window of 20 trials within a single session.

#### 2.2.6 Reversal Learning Phase (REV)

The reversal phase began the day following successful completion of the acquisition phase. In the reversal phase, the apertures designated as correct and incorrect were swapped, but all other contingencies remained the same as those in the acquisition phase – including the performance criterion. Testing continued until criterion was met.

### 2.3 Behavioral Measures and Dependent Variables

#### 2.3.1 Learning Rate

The total number of trials required to reach performance criteria, and the pattern of correct and incorrect choices made over the progression to criteria, were evaluated to characterize individual learning behavior as a function of testing phase. Additionally, to characterize within-phase performance changes, each phase was bisected on the trial in which the maximum change in learning occurred as determined by calculating the maximum change in the cumulative correct response curve (Gallistel et al., 2004; Klanker et al., 2015); leading to a PRE and POST change-point fraction of trials for each testing phase.

#### 2.3.2 Choice Accuracy by Prior Choice (Behavioral Flexibility Score)

Choice behavior was evaluated as a function of feedback from the preceding trial (reward vs timeout punishment) to quantify the tradeoff between behavioral flexibility (changing response after punishment; “Lose-Shift”) and stability (repeating response after reward; “Win-Stay”). Thus, we calculated the proportion of “Lose” and “Win” trials that were correct (p(Lose-Shift) and p(Win-Stay), respectively), then we calculated a flexibility-stability difference score (FS, where FS = p(Lose-Shift) – p(Win-Stay)). Thus, a positive FS would indicate greater behavioral flexibility than stability. Moreover, baseline differences in overall error rates due to other factors are subtracted out by this tradeoff calculation.

#### 2.3.3 Error Parameters (Perseverative and Regressive Errors)

In addition to error rate (errors per trial), two measures parameterized the shape of the learning curve which, in the reversal phase, was characterized by an initial period of perseveration on the old rule followed by a period characterized by occasional “regressive” errors. For the 1^st^ measure, we calculated the maximum length of the strings of consecutive incorrect responses (**MAXCI**) as an operational definition of perseverative responding. For the 2^nd^, we calculated the proportion of the learning curve marked by *regression* from – rather than progression toward – the performance criterion. Learning curves often show variability in these transient periods of decreased proficiency (Gallistel et al., 2004). This **Regress Score** was calculated by identifying blocks of 5 trials in which performance accuracy declined relative to the preceding block, summing the values associated with these declines in accuracy [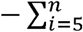 (% correct next 5 trials - % correct prior 5 trials)*i*], then normalizing this sum to the total number of trials.

#### 2.3.4 Premature Responses

Premature responses (nose pokes into the unlit target apertures outside of the period of target presentation) and the number of timeouts/trial incurred from observing response fails (i.e., failures to sustain the observing response until the target stimuli are presented) were parsed as a function of testing **Phase** (acquisition vs reversal; ACQ vs REV), and the within-phase interval relative to the maximum **Change Point** in the learning curve (PRE vs POST). Premature responses were also analyzed with respect to response **Side** (correct side in acquisition or correct side in reversal; **C_ACQ_** or **C_REV_**).

#### 2.3.5 Response Latencies

Target-response latency was the time from target onset until the start of the 1^st^ nose-poke into an illuminated target aperture. Reward-retrieval latency was the time from pellet delivery to the start of 1^st^ subsequent nose-poke into the magazine. Trial-initiation latency was the time from the end of inter-trial interval to the 1^st^ observing response (regardless of whether it was long enough to trigger target presentation).

All three latency measures were analyzed as a function of Testing **Phase** (acquisition vs reversal; ACQ vs REV), and the within-phase interval relative to the maximum **Change Point** in the learning curve (PRE vs POST). Target-Response latencies were parsed by response **Side** (correct side in acquisition or correct side in reversal; **C_ACQ_** or **C_REV_**). Trial-initiation latencies were additionally analyzed by the outcome of the **Prior Trial** (**rewarded** or **unrewarded**).

#### 2.3.6 Extraneous Observing Responses (EOR)

During target-stimuli presentation, additional nose-pokes into the center aperture were recorded but were without scheduled consequences. We analyzed EOR as a function of testing phase and change point.

#### 2.3.7 Omissions

Target stimuli were presented for 30 s. Failure to respond to one of the target apertures within this interval caused a timeout and an omission to be recorded. We analyzed omissions per trial as a function of testing phase and change point.

### 2.4 Statistical Design

Because one of our central hypotheses focused on gonadal males with one vs two X chromosomes, we directly compared the performance of gonadally intact XY vs XXY mice. Separately, we compared XO vs XX mice as a secondary test of the effects of X chromosome number in the presence of ovaries. Measures were analyzed under designs that included the between-subjects factors of X-chromosome number (i.e., **X-dose; 1X** vs **2X**) and gonadectomy group (**GDX**; **GDX** vs **SHAM**). We also analyzed within-subjects factors of testing **Phase** (acquisition vs reversal; **ACQ** vs **REV**), **Prior-Trial Outcome** (**rewarded** vs **unrewarded**), and before/after the maximum change in the learning curve (i.e., **Change Point**; **PRE** vs **POST**; determined by calculating the maximum change in the cumulative correct response curve (Gallistel et al., 2004; Klanker et al., 2015)). Premature responses and target-response latencies were analyzed with reference to response **Side** (correct side in ACQ or correct side in REV; **C_ACQ_** or **C_REV_**).

### 2.5 Data Analyses

Measures were analyzed by the Generalized Estimating Equations (GEE) procedure available in IBM SPSS Statistics version 25 using the robust estimator of the unstructured covariance matrix, maximum likelihood parameter estimation, Type III model effects, and the Chi-square Wald score for full log quasi-likelihood function. A normal distribution with identity link function was used except when the assumptions of normality and linearity were violated (normality determined by Kolmogorov-Smirnov Test (KST), linearity determined by examination of P-P plots). In those cases, a standard data transformation (link function in GEE; options included untransformed, log, or square root) was chosen based on the lowest Chi Square/highest p-value of KSTs of transformed data. Two measures – premature responses/trial, and omissions/trial – could not be normalized by these standard transformations. However, they both adequately fitted, and were thus analyzed using, a gamma distribution with log-link. Significant model effects (p < 0.05) comprising more than two means were delineated with post-hoc paired-means comparisons (PPC) under a Bonferroni correction. Figures plot the raw (untransformed) data, with the exception of premature responses, which were plotted as the value of log_10_(y_i_+1).

## 3 RESULTS

### 3.1 Trials to Criterion

As a measure of learning rate, we counted the total number of response trials required to reach the preset performance criterion (omissions were not counted). Before counting, trials were first parsed by testing phase (acquisition vs reversal: ACQ vs REV) and by a derived intra-phase change point (CP) in the learning curve (PRE vs POST) (Figures 1A and 1B, Supplementary Figures 1A and 1B).

**Figure 1:**
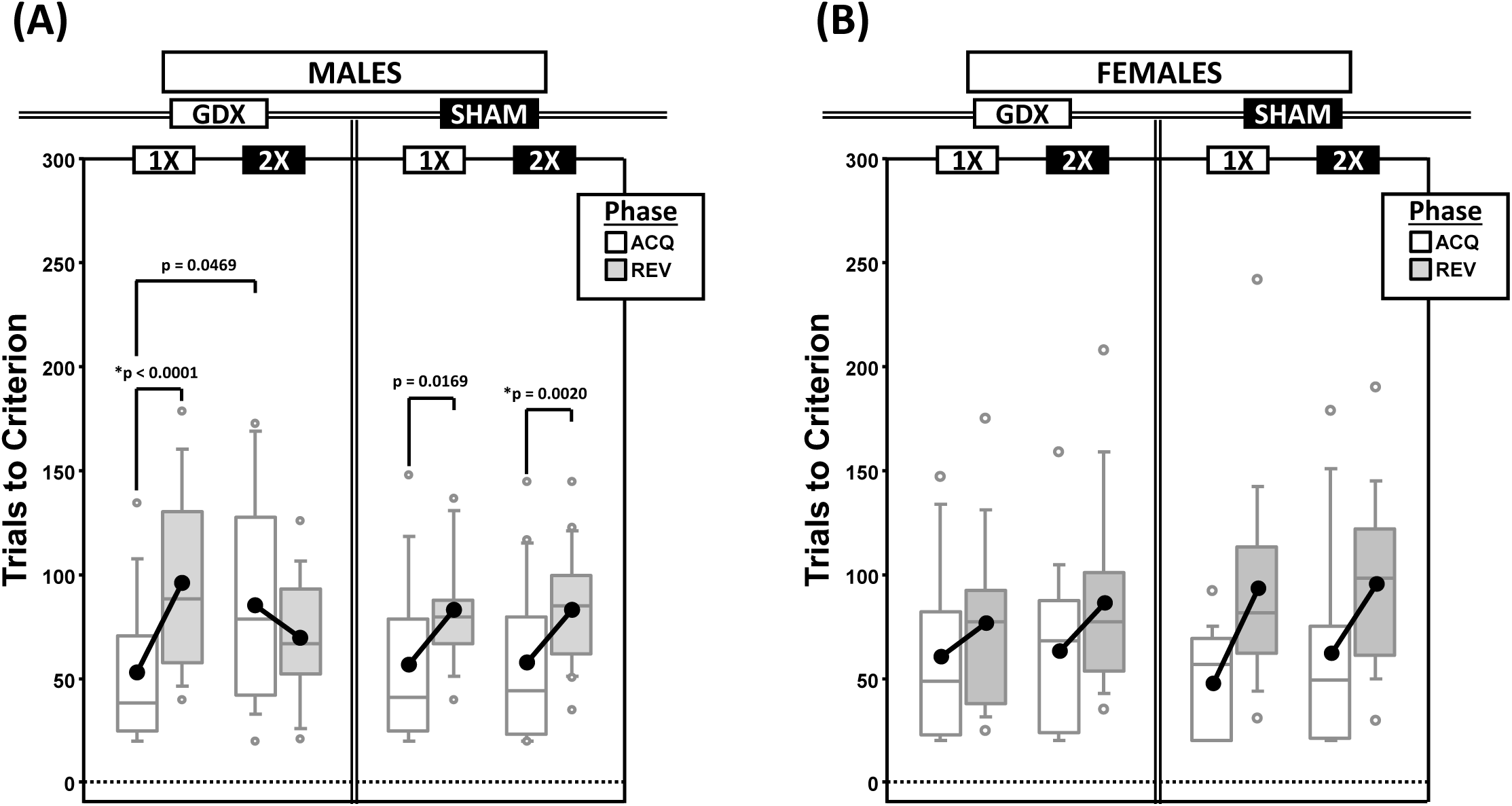
Boxplots of trials to criterion for males (**A**) and for females (**B**) grouped by X-dose (number of X-chromosomes; 1X or 2X) and GDX group, and split by the testing phase (Acquisition = ACQ vs Reversal = REV). Collapsing across and/or splitting by factors in males was based on the omnibus results. Boxes represent median ±quartile, whiskers extend additional 15^th^ of a percentile, grey open circles represent extreme values; black filled circles represent means. Simple effects (p < 0.05) that remained statistically significant after Bonferroni correction are indicated by “*”.

#### 3.1.1 Males

Generalized estimating equations (GEE) analysis confirmed significant effects of Phase (χ^2^_(1)_= 15.935, p < 0.0001) and CP (χ^2^_(1)_= 43.899, p < 0.00001), but there was no Phase*CP interaction (χ^2^_(1)_ = 0.000, p = 0.997). As expected, more trials were required in REV than ACQ and there were more trials PRE than POST (Supplementary Figure 1A).

There were two significant interactions involving experimental groups (Figure 1A): X-dose*Phase (χ^2^_(1)_= 7.938, p = 0.0048) and X-dose*GDX*Phase (χ^2^_(1)_= 7.856, p = 0.0051). Post-hoc paired-means comparisons confirmed a significant simple effect of Phase in both XY-GDX mice (p < 0.0001) and XXY-SHAM mice (p = 0.0020), but the simple effect of Phase in both XY-SHAM mice (p = 0.0169) and XXY-GDX mice (p = 0.18), as well as the simple effect of X-dose (XXY > XY) in GDX mice in ACQ (p = 0.0469) did not survive Bonferroni correction (p > 0.06 for all other simple effects). Thus, although XY-GDX, XY-SHAM, and XXY-SHAM groups showed the expected reversal cost on trials to criterion, the XXY-GDX mice required *fewer* trials in REV than ACQ due to both relatively more trials in ACQ and fewer trials in REV as compared to the other groups, likely reflecting impaired acquisition of the initial rule in this group.

#### 3.1.2 Females

GEE analysis confirmed effects of Phase (χ^2^_(1)_= 21.064, p < 0.00001) and CP (χ^2^_(1)_ = 16.384, p < 0.0001), as well as a Phase*CP interaction (χ^2^_(1)_ = 4.188, p = 0.0407). Post hoc comparisons confirmed that the effect of Phase was significant both PRE (p < 0.00001) and POST (p = 0.005). However, the effect of CP was significant for REV (p = 0.0001), but not for ACQ (p = 0.152). As expected, there were more trials PRE than POST, and more in REV than ACQ (Supplementary Figure 1B). GEE analysis did not confirm any significant effects or interactions involving X-dose or GDX (all p > 0.1) (Figure 1B).

### 3.2 Error Rate

Errors per trial were parsed by testing phase (acquisition vs reversal; ACQ vs REV) and the intra-phase CP in the learning curve (PRE vs POST), the latter to determine within-phase dynamics of learning (Supplementary Figures 1C and 1D).

#### 3.2.1 Males

GEE analysis confirmed significant effects of Phase (χ^2^_(1)_ = 45.172, p < 0.000001) and CP (χ^2^_(1)_ = 551.751, p < 0.000001), as well as a Phase*CP interaction (χ^2^_(1)_ = 18.932, p = 0.00001). As expected, there were more errors made PRE than POST, regardless of phase, and more errors in REV than ACQ, regardless of CP (all p < 0.001; Supplementary Figure 1C). There were no significant effects or interactions involving X-dose or GDX (all p > 0.1).

#### 3.2.2 Females

GEE analysis confirmed effects of Phase (χ^2^_(1)_ = 23.670, p < 0.00001) and CP (χ^2^_(1)_ = 610.346, p < 0.00001), and a Phase*CP interaction (χ^2^_(1)_ = 6.873, p = 0.0088). As expected, there were more errors made PRE than POST, regardless of phase, and more errors in REV than ACQ, regardless of CP (all p < 0.0001; Supplementary Figure 1D). There were no significant effects or interactions involving X-dose or GDX (all p > 0.1).

### 3.3 Behavioral Flexibility Score (FS)

To score the tradeoff between behavioral flexibility and stability, we calculated the difference between the proportion of responses that were correct following punished trials (p(Lose-Shift); index of flexibility) vs those that were correct following rewarded trials (p(Win-Stay); index of stability) (FS = p(Lose-Shift) – p(Win-Stay); Supplementary Figures 2A and 2B). Thus, a positive FS score would indicate greater behavioral flexibility than stability. These calculations were based on observations parsed by both the testing phase (ACQ vs REV) and the CP in the learning curve (PRE vs POST; used to examine within-phase dynamics).

#### 3.3.1 Males

As expected, mean FS decreased from a positive value PRE to a negative value POST in both ACQ and REV (Supplementary Figure 2A), reflecting a shift from a Lose-Shift strategy to a Win-Stay strategy. GEE analysis confirmed significant main effects of CP (χ^2^_(1)_ = 38.214, p < 0.00001) and Phase (χ^2^_(1)_ = 9.825, p = 0.0017), but no Phase*CP interaction (χ^2^_(1)_ = 0.019, p = 0.89). Additionally, the overall mean FS was positive in SHAM mice but negative in GDX mice, confirmed by a main effect of GDX (χ^2^_(1)_ = 5.088, p = 0.024) (Figure 2A).

**Figure 2:**
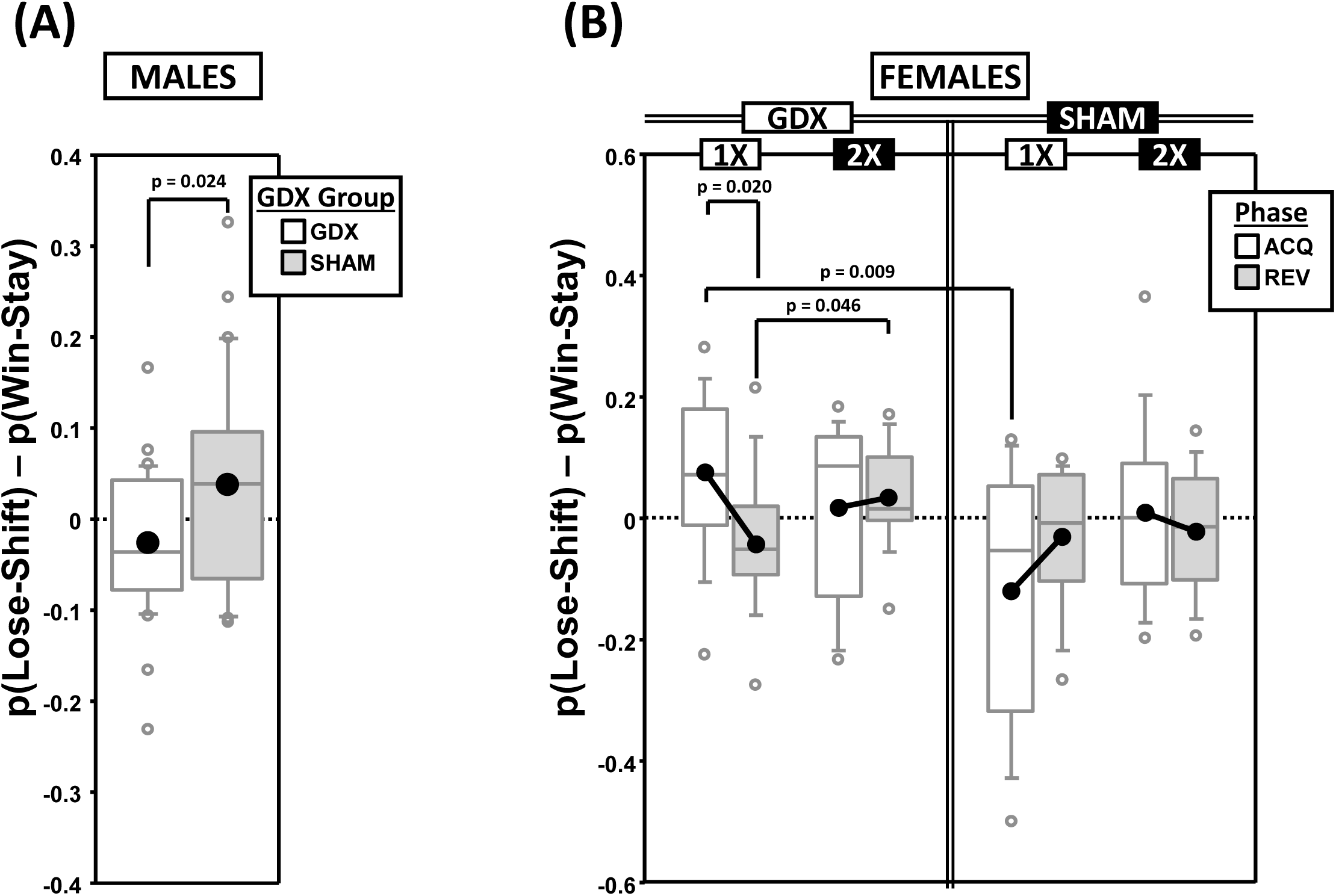
Boxplots of the difference between the proportion of responses that were correct following either an incorrect response (**p(Lose-Shift)**; index of *flexibility*) or a correct response (**p(Win-Stay)**; index of *stability*) for males (**A**) broken down by gonadectomy group, or for females (**B**) broken down by X-dose (number of X-chromosomes; 1X or 2X), gonadectomy group and testing phase. Collapsing across and/or splitting by factors was based on the omnibus results. Boxes represent median ±quartile, whiskers extend additional 15^th^ of a percentile, grey open circles represent extreme values; black filled circles represent means. None of the simple effects (p < 0.05) indicated in (B) were significant after Bonferroni correction.

#### 3.3.2 Females

GEE analysis of FS confirmed a significant main effect of CP (χ^2^_(1)_ = 27.965, p < 0.000001), but not of Phase (χ^2^_(1)_ = 0.732, p = 0.39) nor a Phase*CP interaction (χ^2^_(1)_ = 1.943, p = 0.16), reflecting the fact that FS decreased from a positive value PRE to a negative value POST in both ACQ and REV (Supplementary Figure 2B). GEE also confirmed an X-dose*GDX*Phase interaction (χ^2^_(1)_ = 4.821, p = 0.028) (Figure 2B). However, the simple effects of X-dose (XX > XO in GDX in REV; p = 0.046), of GDX (GDX > SHAM in XO in ACQ; p = 0.009), and of Phase (ACQ > REV in GDX XO; p = 0.020) did not survive correction for multiple comparisons.

### 3.4 Maximum Number of Consecutive Incorrect Responses (MAXCI)

To quantify response perseveration, we calculated the MAXCI within each testing phase (ACQ scores served as a baseline from which the magnitude of the effect of reversal on perseveration was derived) (Figures 3A and 3B). This was an alternative way of quantifying perseveration as compared to methods based on the number of errors within sequential blocks of trials (e.g., (Ragozzino et al., 2002)). The MAXCI was chosen for two reasons: 1) Prior analysis of cumulative response records from this particular reversal task indicated that it is more sensitive to the magnitude of the initial bout of post-reversal perseveration and 2) it consistently shows no significant correlation with the complementary regressive error measure (Figure 3D).

**Figure 3:**
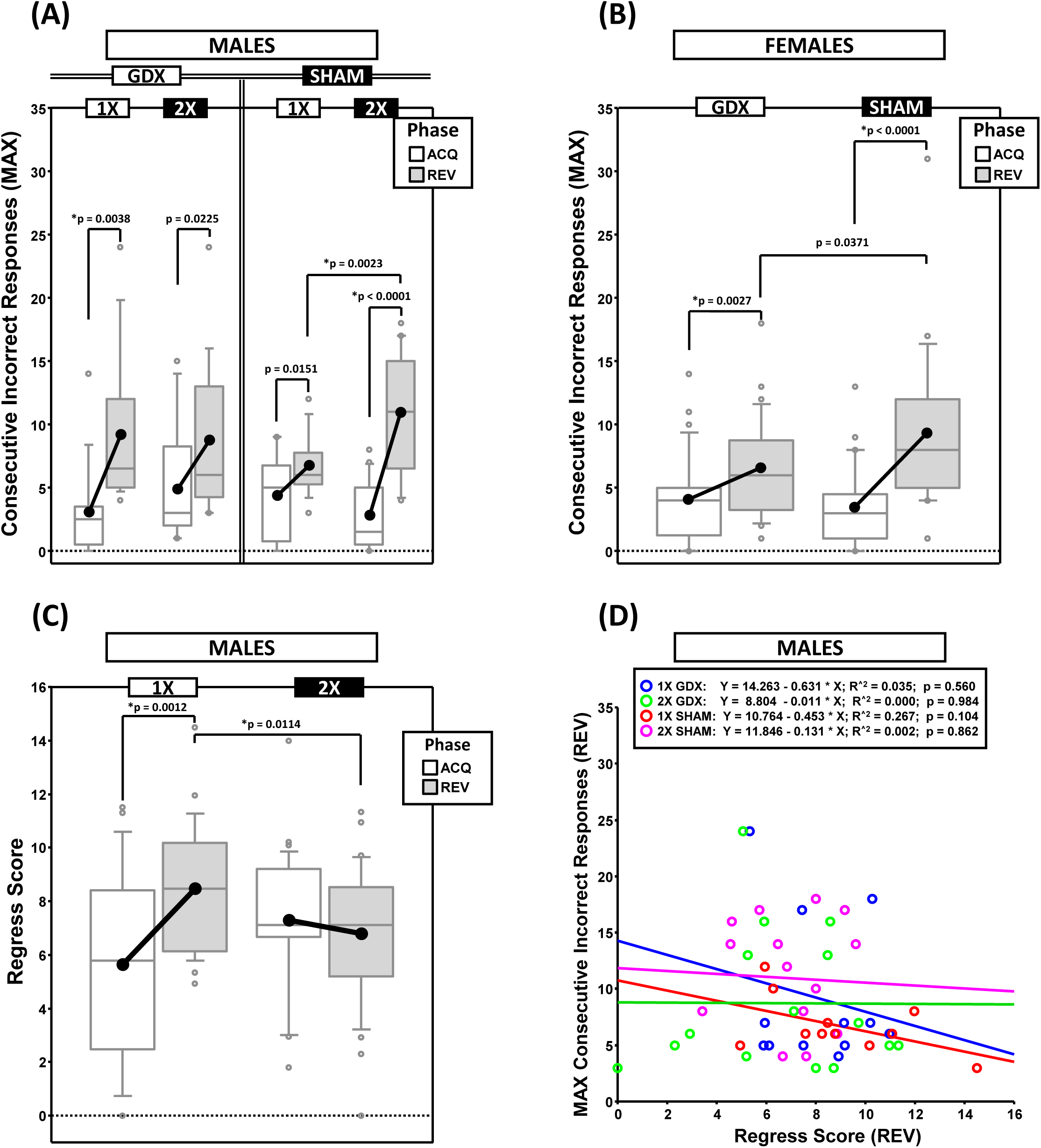
Boxplots of maximum consecutive incorrect responses for males (**A**) and for females (**B**), as well as the regress score for males (**C**) broken down by testing phase (Acquisition = ACQ vs Reversal = REV). Collapsing across and/or splitting by factors was based on the omnibus results. Boxes represent median ±quartile, whiskers extend additional 15^th^ of a percentile, grey open circles represent extreme values; black filled circles represent means. Simple effects (p < 0.05) on either measure that remained statistically significant after Bonferroni correction are indicated by “*”. (**D**) Bivariate scattergram of maximum consecutive incorrect and regress score for males split by group.

#### 3.4.1 Males

GEE confirmed a main effect of Phase (χ^2^_(1)_ = 45.719, p < 0.00001) and a significant X-dose*GDX*Phase interaction (χ^2^_(1)_ = 7.189, p = 0.0073) (Figure 3A). Post hoc comparisons confirmed a simple effect of X-dose in the SHAM group in REV (XXY > XY; p = 0.0023; all other p > 0.19). By contrast, no simple effects of GDX were confirmed (all p > 0.13). Lastly, although MAXCI was greater in REV than ACQ for XY-GDX (p = 0.0038), XY-SHAM, (p = 0.0151), XXY-GDX (p = 0.0225), and XXY-SHAM (p < 0.00001), this difference was not significant for XY-SHAM and XXY-GDX after Bonferroni correction.

#### 3.4.2 Females

GEE analysis confirmed a main effect of Phase (χ^2^_(1)_ = 36.010, p < 0.00001) and a significant GDX*Phase interaction (χ^2^_(1)_ = 4.924, p = 0.027). Although the simple effect of Phase in GDX (REV > ACQ; p = 0.0027) and SHAM (REV > ACQ; p < 0.0001) survived correction for multiple comparisons, the simple effect of GDX in REV did not (SHAM > GDX; p = 0.0371) (Figure 3B).

### 3.5 Regress Score (regressive errors on path to criterion)

To quantify the tendency for performance to initially improve and then regress away from criterion, we calculated the changes in accuracy across the phase (moving window: prior 5 trials – next 5 trials), summed the negative values and then normalized this sum to the total number of trials experienced (Figure 3C).

#### 3.5.1 Males

GEE analysis confirmed a main effect of Phase (χ^2^_(1)_ = 4.792, p = 0.0286), and an X-dose*Phase interaction (χ^2^_(1)_ = 10.348, p = 0.0013) (Figure 3C). Post hoc comparisons confirmed that Regress scores were higher in REV than ACQ in XY mice (p = 0.0012), but not XXY mice (p = 0.35). Additionally, the Regress score was higher in XY mice than XXY mice in REV (p = 0.0114), but not in ACQ (p = 0.0517).

#### 3.5.2 Females

GEE analysis did not confirm any main effects or interactions (all p > 0.14).

### 3.6 Correlations: MAXCI, Regress Score, and Trials to Criterion

If MAXCI and the Regress Score quantify independent components of reversal learning (i.e., perseverative and regressive errors, respectively), then they should not be correlated.

Correlational analyses split by sex, X-dose and GDX groups confirmed that these measures were not correlated (all p > 0.1, males illustrated in Figure 3D). However, both MAXCI and the Regress Score were positively correlated to trials to criterion (MAXCI, all p < 0.008; Regress Score, all p < 0.007). Thus, although errors of either type predicted trials to criterion, one error did not predict the other.

### 3.7 Premature Responding

Premature responses per trial to the target apertures (i.e., nose pokes into the unlit target apertures outside of the period of target presentation) on both the correct side in acquisition (**C_ACQ_**) and the correct side in reversal (**C_REV_**) were analyzed as a function of testing phase and change point (CP) (Figures 4A thru 4G).

**Figure 4:**
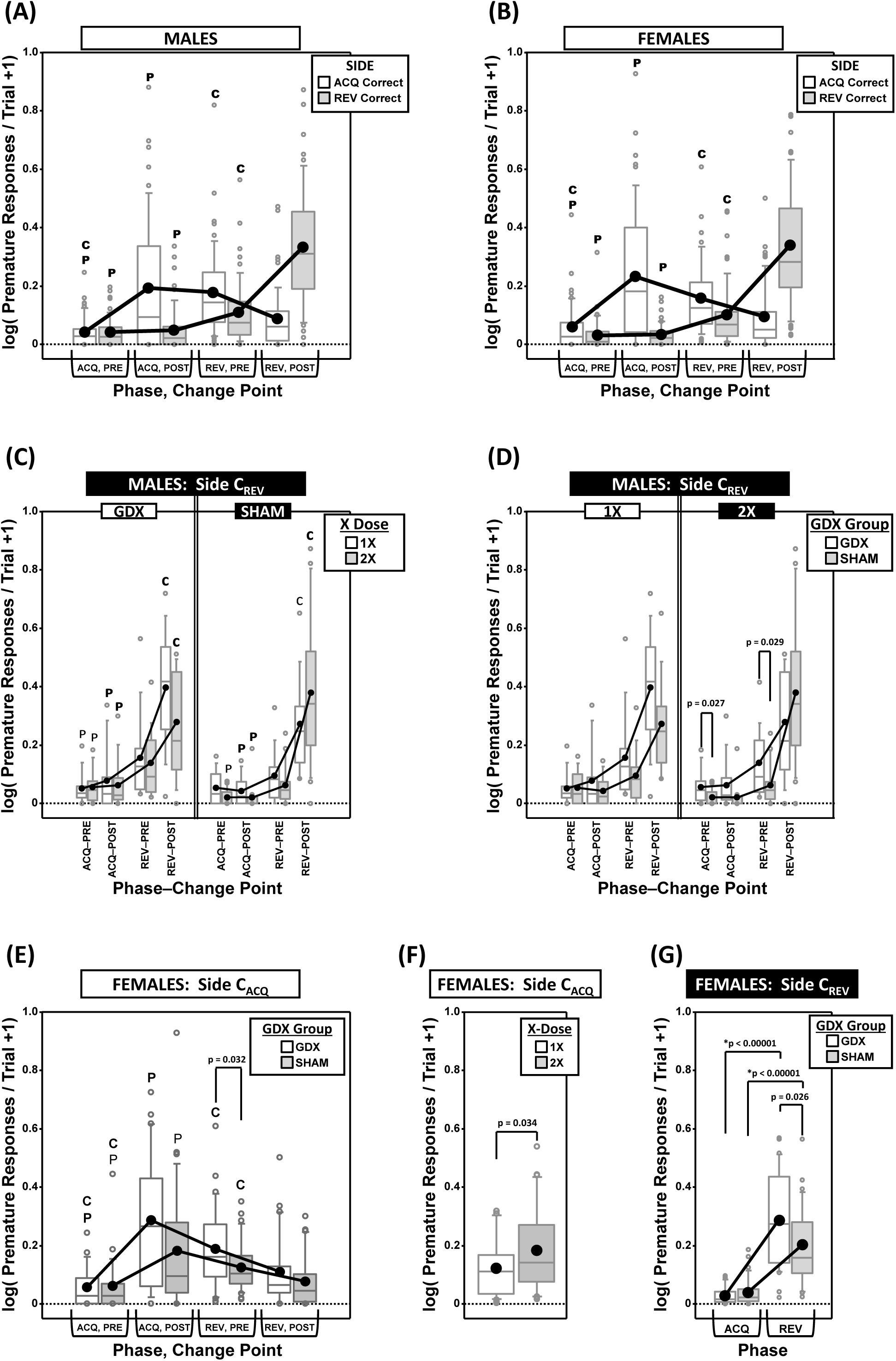
Top panels are boxplots of log_10_ of premature responses per trial (+1) split by response side (correct in ACQ vs correct in REV; **C_ACQ_** vs **C_REV_**), parsed by testing phase (Acquisition = ACQ vs Reversal = REV) and Intra-phase Change Point (PRE vs POST) for males **(A)** and for females **(B)**. Middle panels are premature responses for males on the **C_REV_** side that are either split by X-dose (number of X-chromosomes; 1X or 2X) and grouped by GDX group **(C)**, or vice versa **(D)**. Bottom panels are premature responses for females on the **C_ACQ_** side that are either split by GDX group and parsed by testing phase and change point **(E),** or only split by X-dose **(F)**, and premature responses for females on the **C_REV_** side split by GDX group and parsed by testing phase **(G)**. Collapsing across and/or splitting by factors was based on the omnibus results. Simple effects of group (p < 0.05) that remained statistically significant after Bonferroni correction are indicated by “*”. Letters “P” and “C” indicate simple effects of Phase and Change Point (p < 0.05), respectively (bold letters indicate significance after Bonferroni correction). Boxes represent median ±quartile, whiskers extend additional 15^th^ of a percentile, grey open circles represent extreme values; black filled circles represent means.

#### 3.7.1 Males

GEE analysis of premature responding on the **C_ACQ_** side confirmed a main effect of CP (χ^2^_(1)_ = 7.905, p = 0.0049) and a Phase*CP interaction (χ^2^_(1)_ = 41.553, p < 0.00001), but no main effect of Phase (χ^2^_(1)_ = 0.008, p = 0.93) (Figure 4A). However, neither X-dose, nor GDX, affected this pattern of results (all p > 0.15). The simple effects of Phase and Change point were all significant (all p < 0.001).

GEE analysis of premature responding on the **C_REV_** side confirmed main effects of Phase (χ^2^_(1)_ = 154.663, p < 0.00001), CP (χ^2^_(1)_ = 72.122, p < 0.00001), and a Phase*CP interaction (χ^2^_(1)_ = 72.995, p < 0.00001). There was also an X-dose*GDX*Phase interaction (χ^2^_(1)_ = 6.067, p = 0.0138), X-dose*GDX*CP interaction (χ^2^_(1)_ = 6.958, p = 0.0083), X-dose*GDX*Phase*CP interaction (χ^2^_(1)_ = 4.488, p = 0.034), and a GDX*Phase*CP interaction (χ^2^_(1)_ = 4.752, p = 0.029) (Figures 4C and 4D). There were no significant simple effects of X-dose (all p > 0.05) and the simple effects of GDX (GDX > SHAM in XXY in both ACQ PRE and REV PRE; p = 0.027 and p = 0.029, respectively) did not survive correction for multiple comparisons (Figure 4D) (see Figure 4C for simple effects of Phase and CP).

#### 3.7.2 Females

GEE analysis of premature responding on the **C_ACQ_** side confirmed a main effect of CP (χ^2^_(1)_ = 31.629, p < 0.00001) and a Phase*CP interaction (χ^2^_(1)_ = 77.525, p < 0.00001), but no main effect of Phase (χ^2^_(1)_ = 2.378, p = 0.12) (Figure 4B; all simple effects of Phase and Change point significant; all p < 0.0001). GEE analysis also confirmed a main effect of X-dose (Figure 4C; χ^2^_(1)_ = 4.506, p = 0.034), with XX mice overall exhibiting more premature responses than XO subjects (Figure 4F). Finally, GEE confirmed a GDX*Phase*CP interaction (χ^2^_(1)_ = 4.084, p= 0.043), but none of the post hoc comparisons for the simple effect of GDX were significant after correction (all p > 0.03) (see Figure 4E for simple effects of Phase and CP).

GEE analysis of premature responding on the **C_REV_** side confirmed main effects of Phase (χ^2^_(1)_ = 160.374, p < 0.00001) and CP (χ^2^_(1)_ = 88.466, p < 0.00001), as well as a Phase*CP interaction (χ^2^_(1)_ = 86.810, p < 0.00001; see Figure 4B for simple effects) and a GDX*Phase interaction (χ^2^_(1)_ = 7.773, p = 0.0053; Figure 4G). Although post hoc comparisons confirmed the simple effect of phase within both GDX groups (both p < 0.00001), the simple effect of GDX in REV (GDX > SHAM, p = 0.026) did not survive Bonferroni correction.

### 3.8 Response Latencies

In addition to a decrease in response errors, successful learning of a spatial discrimination task leads to progressive decreases in response latencies. Thus, we calculated mean response latencies (i.e., time from target stimuli onset to first response to a target location) on both the correct side in acquisition (**C_ACQ_**) and the correct side in reversal (**C_REV_**) and analyzed these as a function of testing phase and CP (Figures 5A and 5B, Supplementary Figures 3A and 3B).

**Figure 5:**
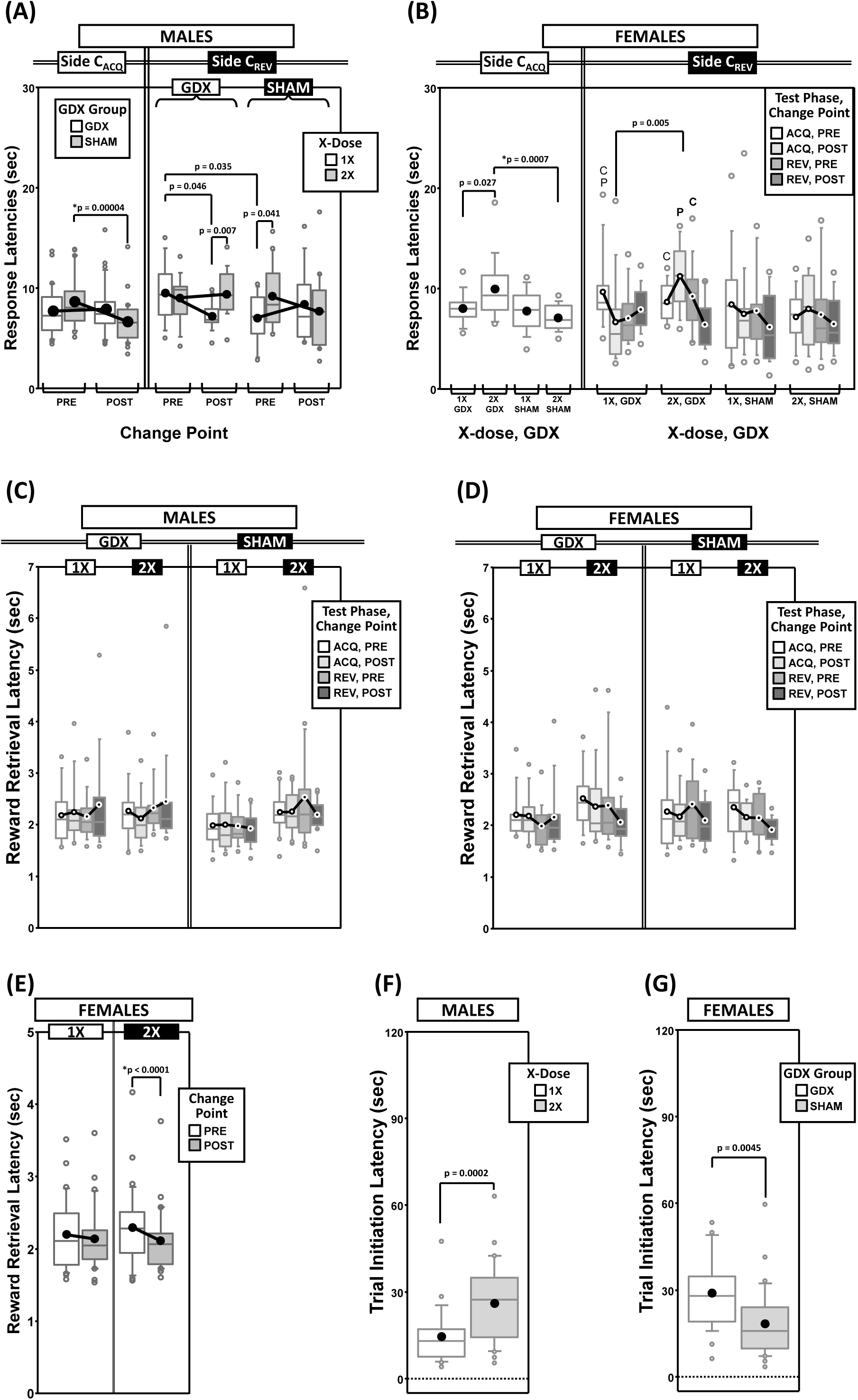
Top panels are boxplots of response latencies to target apertures as a function of response side (correct in ACQ = **C_ACQ_** vs correct in REV = **C_REV_**) for males **(A)** and for females **(B)**, split by X-dose (number of X-chromosomes; 1X or 2X) or GDX group and parsed by testing phase (Acquisition = ACQ vs. Reversal = REV) and/or Intra-phase Change Point (PRE vs. POST). Middle panels are boxplots of reward-retrieval latencies for males (**C**) and for females (**D**) grouped by X-dose, GDX group, testing phase and CP. Bottom panels include box plots of reward retrieval latencies for females collapsed across testing phase and GDX group **(E)** and of trial-initiation latencies for males split by X-dose (**F**) and for females split by GDX group (**G**). Collapsing across and/or splitting by factors was guided by the omnibus results. Simple effects of group (p < 0.05) on response latencies and trial initiation latencies that remained statistically significant after Bonferroni correction are indicated by “*”. Letters “P” and “C” indicate simple effects of Phase and Change Point (p < 0.05), respectively, on response latencies and trial initiation latencies (bold letters indicate significance after Bonferroni correction). On reward retrieval latencies in males, there was a significant GDX by Phase by Change point interaction (p = 0.048; no significant simple effects, all p > 0.06). On reward retrieval latencies in females, there were main effects of Phase (p = 0.042) and Change Point (p = 0.011), and an X-dose by Change Point interaction (p = 0.033; simple effect of Change Point in 2X, p < 0.0001; all other simple effects, p > 0.12). Boxes represent median ±quartile, whiskers extend additional 15^th^ of a percentile, grey open circles represent extreme values; black filled circles represent means.

#### 3.8.1 Males

GEE analysis of response latencies on the **C_ACQ_** side confirmed main effects of Phase (χ^2^_(1)_ = 16.226, p < 0.0001) and CP (χ^2^_(1)_ = 7.357, p = 0.007), as well as a Phase*CP interaction (χ^2^_(1)_ = 9.700, p = 0.002). However, on the **C_REV_** side, analysis confirmed a main effect of Phase (χ^2^_(1)_ = 18.521, p = 0.00002), but no effect of CP (χ^2^_(1)_ = 0.990, p = 0.3) or Phase*CP interaction (χ^2^_(1)_ = 1.465, p = 0.2). Response latencies to the **C_ACQ_** side decreased during ACQ (POST CP < PRE CP) but not REV (POST CP ≈ PRE CP), while response latencies to the **C_REV_** side decreased during REV (POST CP < PRE CP) but not ACQ (POST CP ≈ PRE CP) (Supplementary Figure 3A).

Analysis of the **C_ACQ_** side also confirmed a significant GDX*CP interaction (χ^2^_(1)_ = 11.732, p = 0.0006). Post hoc comparisons confirmed a simple effect of CP in SHAM mice (p = 0.00004) but not in GDX mice (p = 0.6); the simple effect of GDX was not significant either PRE (p = 0.14) or POST (p = 0.07) (Figure 5A).

Lastly, an X-dose*GDX*CP interaction was confirmed on the **C_REV_** side (χ^2^_(1)_ = 6.130, p = 0.013), but post hoc comparisons did not identify any simple effects that survived correction for multiple comparisons (Figure 5A).

#### 3.8.2 Females

GEE analysis of response latencies on the **C_ACQ_** side confirmed main effects of Phase (χ^2^_(1)_ = 22.846, p < 0.00001) and CP (χ^2^_(1)_ = 5.234, p = 0.02), but no Phase*CP interaction (χ^2^_(1)_ = 0.651, p = 0.4) (Supplementary Figure 3B). A main effect of GDX (χ^2^_(1)_ = 8.007, p = 0.005), as well as an X-dose*GDX interaction (χ^2^_(1)_ = 5.791, p = 0.016) was identified (Figure 5B).

Post hoc comparisons confirmed the simple effect of GDX in XX mice (p = 0.0007); however the simple effect of X-dose in GDX mice was not significant after Bonferroni correction (p = 0.027) (other p > 0.2).

GEE analysis of response latencies on the **C_REV_** side confirmed a main effect of Phase (χ^2^_(1)_ = 5.996, p = 0.014), but not of CP (χ^2^_(1)_ = 3.019, p = 0.08) nor a Phase*CP interaction (χ^2^_(1)_ = 1.503, p = 0.2) (Supplementary Figure 3B). GEE analysis also confirmed an X-dose*Phase*CP interaction (χ^2^ = 8.276, p = 0.004) and an X-dose*GDX*Phase*CP interaction (χ^2^ = 4.361, p = 0.037). No simple effects of X-dose or GDX survived correction for multiple comparisons (Figure 5B).

### 3.9 Reward Retrieval Latencies

As an ancillary response latency measure, we calculated the average time to collect the reward after committing a correct response as a function of both testing phase and change point (CP) (Figures 5C thru 5D).

#### 3.9.1 Males

GEE analysis of reward retrieval latencies confirmed no main effects of Phase (χ^2^_(1)_ = 0.468, p = 0.5) or CP (χ^2^_(1)_ = 0.130, p = 0.7), nor a Phase*CP interaction (χ^2^_(1)_ = 0.059, p = 0.8). GEE analysis did confirm a GDX*Phase*CP interaction (χ^2^_(1)_ = 3.914, p = 0.048), however none of the post hoc comparisons were significant (all p > 0.06) (Figure 5C).

#### 3.9.2 Females

GEE analysis of reward retrieval latencies did confirm main effects of Phase (χ^2^_(1)_ = 4.151, p = 0.04; ACQ > REV) and CP (χ^2^_(1)_ = 6.495, p = 0.01; PRE > POST), but no Phase*CP interaction (χ^2^_(1)_ = 0.257, p = 0.6) (Figure 5D). GEE analysis also confirmed an X-dose*CP interaction (χ^2^_(1)_ = 4.539, p = 0.03). Post hoc comparisons confirmed a simple effect of CP in XX mice (p < 0.0001) (all other p > 0.1) (Figure 5E).

### 3.10 Trial Initiation Latencies

As an ancillary response latency measure that is affected by motivation to engage in the task, we calculated the mean time to initiate a trial (i.e., time from the end of inter-trial interval to the 1^st^ observing response) as a function of both testing phase and the outcome of the prior trial (rewarded or unrewarded) (Figure 5F and 5G, Supplementary Figures 3C and 3D).

#### 3.10.1 Males

As expected, trial initiation latencies were shorter after unrewarded trials (M = 12.0 s, SD = 8.3 s) than after rewarded trials (M = 29.5 s, SD = 20.1 s), regardless of phase (Supplementary Figure 3C). GEE analysis on latencies confirmed main effects of Phase (χ^2^_(1)_ = 11.387, p = 0.0007) and the outcome of the prior trial (χ^2^_(1)_ = 134.876, p < 0.00001), but no Phase*Prior-Trial interaction (χ^2^_(1)_ = 1.473, p = 0.2). GEE also confirmed a main effect of X-dose (χ^2^_(1)_ = 14.300, p = 0.00016). XXY mice had overall longer trial initiation latencies than XY mice (Figure 5F).

#### 3.10.2 Females

As expected, trial initiation latencies were shorter after unrewarded trials (M = 14.4 s, SD = 9.8 s) than rewarded trials (M = 33.3 s, SD = 18.2 s) (Supplementary Figure 3D). GEE analysis on latencies confirmed main effects of Phase (χ^2^_(1)_ = 4.379, p = 0.036) and the outcome of the prior trial (χ^2^_(1)_ = 151.134, p < 0.00001), but no Phase*Prior-Trial interaction (χ^2^_(1)_ = 1.086, p = 0.3). GEE also confirmed a main effect of GDX (χ^2^_(1)_ = 8.088, p = 0.0045). GDX mice had overall longer latencies than SHAM mice (Figure 5G).

### 3.11 Timeouts from Observing-Response Failures (ORTO)

Failure to complete the holding requirement of the observing response (∼0.0 to 0.2 seconds) causes a short 2-second timeout after which the subject must make another attempt; thus, a subject can incur multiple observing-response timeouts each trial. These errors may reflect a facet of response inhibition or impulsivity (Linden et al., 2018); thus, we calculated the mean number of timeouts per trial as a function of both testing phase and change point (CP) (Figures 6A and 6B).

**Figure 6:**
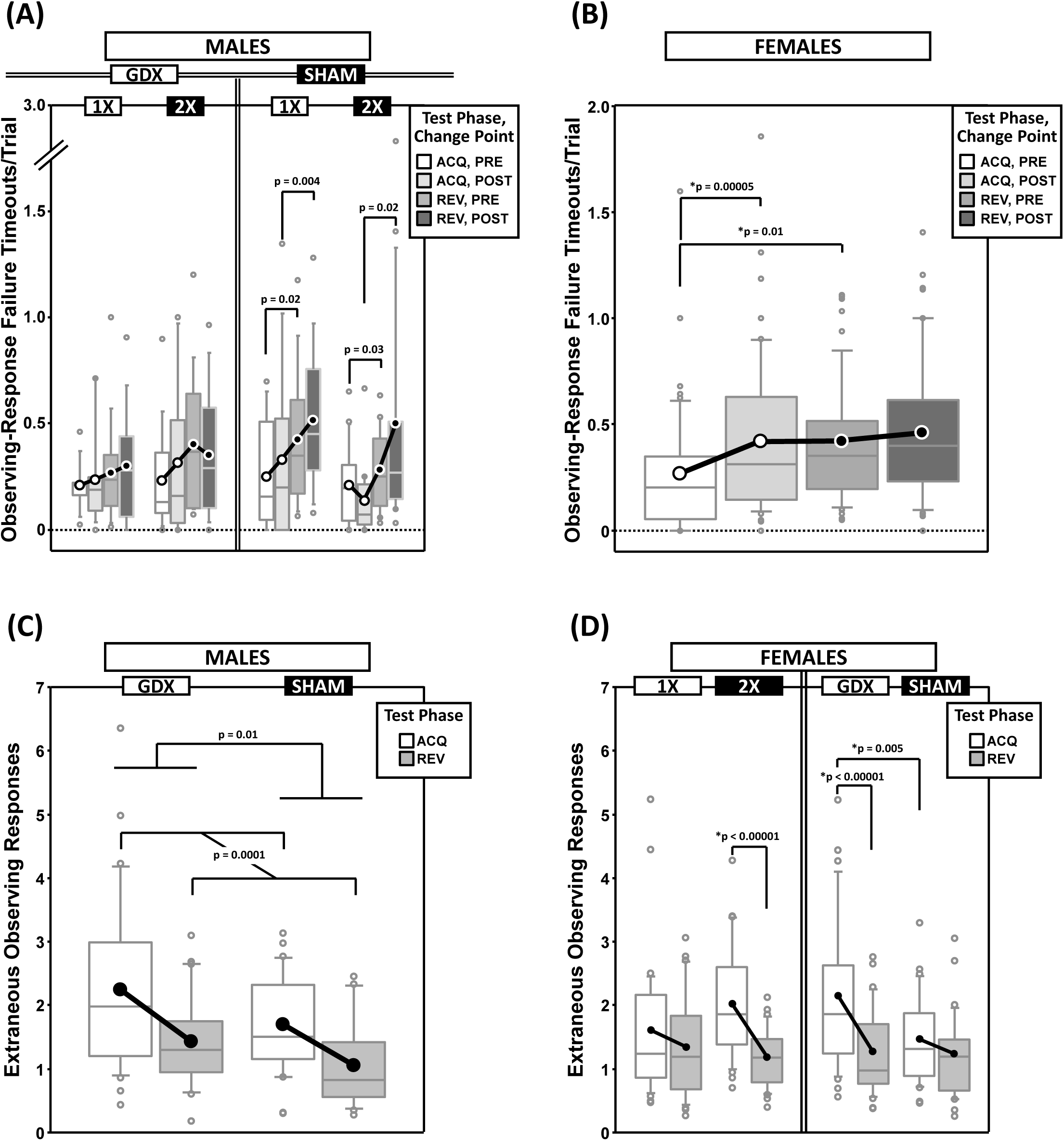
Top panels are boxplots of timeouts/trial due to failures to complete the observing response for males (**A**) and for females (**B**), split by X-dose and/or GDX group, and parsed by testing phase (Acquisition = ACQ vs. Reversal = REV) and Intra-phase Change Point (PRE vs. POST). In males, the main effect of Phase was significant (p = 0.0001). Bottom panels are boxplots of extraneous observing responses/trial (i.e., observing responses made during target presentation) for males grouped by GDX group (**C**) and for females grouped by GDX group or X-dose (**D**) – both parsed by testing phase. In females, the main effect of Phase was significant (p = 0.00003). Collapsing across and/or splitting by factors was based on the omnibus results. Simple effects (p < 0.05) that remained statistically significant after Bonferroni correction are indicated by “*”. Boxes represent median ±quartile, whiskers extend additional 15^th^ of a percentile, grey open circles represent extreme values; black/white circles filled white/black represent means.

#### 3.11.1 Males

GEE analysis of ORTO confirmed a main effect of Phase (REV > ACQ; χ^2^_(1)_ = 18.311, p = 0.0001), but no main effect of CP (χ^2^_(1)_ = 2.089, p = 0.15), nor a Phase*CP interaction (χ^2^_(1)_ = 0.611, p = 0.4). GEE analysis also confirmed a GDX*Phase*CP interaction (χ^2^_(1)_ = 4.263, p = 0.039) and an X-dose*GDX*Phase*CP interaction (χ^2^_(1)_ = 4.780, p = 0.029). However, none of the post hoc comparisons were significant after Bonferroni correction (Figure 6A).

#### 3.11.2 Females

GEE analysis of ORTO confirmed a main effect of CP (χ^2^_(1)_ = 10.091, p = 0.0015) and a Phase*CP interaction (χ^2^_(1)_ = 4.075, p = 0.044), but no main effect of Phase (χ^2^_(1)_ = 3.103, p = 0.078) (Figure 6B). Post hoc comparisons confirmed a simple effect of Phase PRE (REV > ACQ; p = 0.01) and a simple effect of CP in ACQ (POST > PRE; p = 0.00005) (other p > 0.3).

### 3.12 Extraneous Observing Responses (EOR)

During the period of target aperture illumination, observing responses were without consequence. However, as these extraneous observing responses (EOR) may reflect a lack of attention and/or the degree to which the subject has learned the task rules (EOR are expected to decrease and the animal gains proficiency), we calculated the mean EOR per trial parsed by Phase and CP (Figures 6C and 6D).

#### 3.12.1 Males

GEE analysis of EOR confirmed a main effect of Phase (χ^2^_(1)_ = 14.598, p = 0.0001) and GDX (χ^2^_(1)_ = 6.018, p = 0.014), but no main effect of CP (χ^2^_(1)_ = 0.023, p = 0.9) nor Phase*CP interaction (χ^2^_(1)_ = 0.076, p = 0.8). EOR decreased from ACQ to REV and were higher in GDX mice than SHAM mice (Figure 6C).

#### 3.12.2 Females

GEE analysis of EOR confirmed a main effect of Phase (ACQ > REV; χ^2^_(1)_ = 17.410, p = 0.00003) and GDX (χ^2^_(1)_ = 4.647, p = 0.03), but no main effect of CP (χ^2^_(1)_ = 0.724, p = 0.4) nor Phase*CP interaction (χ^2^_(1)_ = 0.971, p = 0.3). GEE analysis also confirmed an X-dose*Phase interaction (χ^2^_(1)_ = 5.013, p = 0.025) and a GDX*Phase interaction (χ^2^_(1)_ = 6.296, p = 0.012). Post hoc comparisons confirmed simple effects of Phase in XX mice (p < 0.00001) and GDX mice (p < 0.00001), as well as a simple effect of GDX in ACQ (GDX > SHAM; p = 0.005) (all other p > 0.08) (Figure 6D).

### 3.13 Omissions

Failure to make a response into a target aperture during the 30-s target-stimuli presentation interval resulted in a 5-s timeout and the recording of an omission error. We calculated the proportion of trials that were omission errors parsed by testing phase and CP (Figures 7A and 7B, Supplementary Figures 4A and 4B).

**Figure 7:**
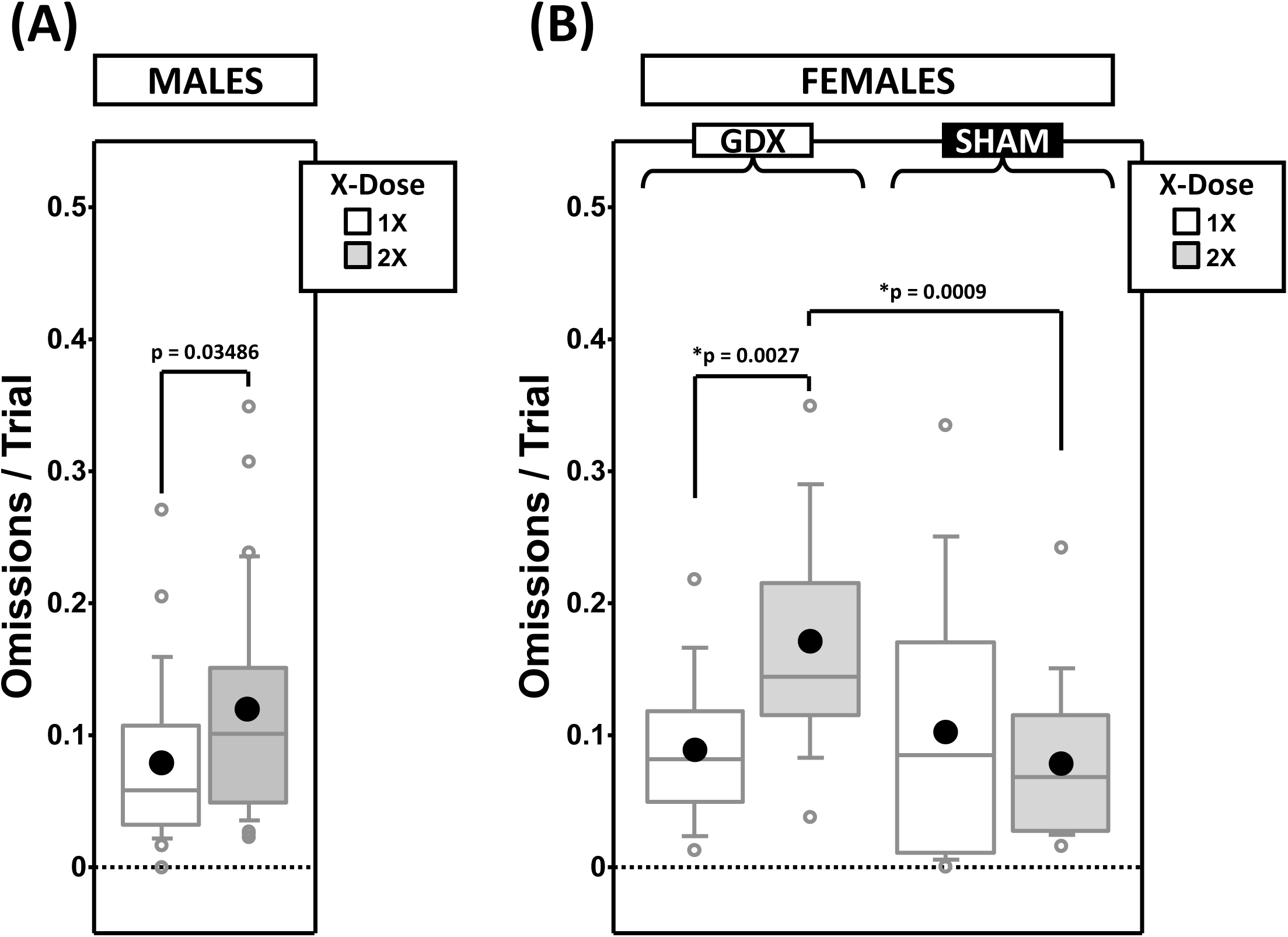
Boxplots of omissions per trial for males split by X-dose **(A)** and for females split by X-dose and GDX group **(B)**. Collapsing across and/or splitting by factors was based on the omnibus results. Simple effects (p < 0.05) that remained statistically significant after Bonferroni correction are indicated by “*”. Boxes represent median ±quartile, whiskers extend additional 15^th^ of a percentile, grey open circles represent extreme values; black filled circles represent means.

#### 3.13.1 Males

GEE analysis of omission errors confirmed a main effect of X-dose (χ^2^_(1)_ = 4.452, p = 0.035), Phase (χ^2^_(1)_ = 16.845, p = 0.00004), and CP (χ^2^_(1)_ = 67.598, p < 0.00001), as well as a Phase*CP interaction (χ^2^_(1)_ = 12.507, p = 0.0004). As expected, omission errors decreased from ACQ-PRE to REV-POST (Supplementary Figure 4A). Omission errors were overall more frequent in XXY than XY mice (Figure 7A).

#### 3.13.2 Females

GEE analysis of omission errors confirmed the main effects of Phase (χ^2^_(1)_ = 10.543, p = 0.001) and CP (χ^2^_(1)_ = 68.951, p < 0.00001), but no Phase*CP interaction (χ^2^_(1)_ = 2.164, p = 0.14). As expected, omission errors decreased from ACQ-PRE to REV-POST (Supplementary Figure 4B). GEE analysis also confirmed an X-dose*GDX interaction (χ^2^_(1)_ = 6.671, p = 0.0098). Post hoc comparisons confirmed simple effects of X-dose in GDX mice (XX > XO; p = 0.0027) and of GDX in XX mice (GDX > SHAM; p = 0.0009) (other p > 0.4). Omissions were overall more frequent in XX-GDX mice than in the other groups (Figure 7B).

## 4 DISCUSSION

### 4.1 Overview

In gonadally-intact males, XXY mice exhibited a greater degree of perseverative error in the reversal phase than XY mice, while in gonadectomized males, XXY mice required more trials to reach criterion in the acquisition phase than XY mice. These results indicate that gonadal function moderates the impact of X-dose on impulsivity and learning, respectively, in the XY* model of KS. Nevertheless, in KS men, androgen therapy in adulthood does not appear to affect these aspects of learning and cognition (Fales et al., 2003; Kompus et al., 2011; Liberato et al., 2017; Skakkebæk et al., 2017).

Additionally, a number of gonadectomy-independent effects were observed (overall, longer trial initiation latencies and more omissions by XXY compared to XY; greater regressive error in reversal in XY than XXY) that indicate direct effects of the sex-chromosome complement on behavior, even if the relevance of these effects to KS is currently either unclear or unsubstantiated.

Consistent with past reports (Davies et al., 2005, 2007), XO vs XX comparisons of RLT performance do not model the cognitive deficits in TS.

### 4.2 XXY vs XY mice as a model of KS

Two principle measures discriminated XXY from XY mice – trials to criterion and perseverative responding in reversal – and both of these interacted with gonadal status. The group difference in perseverative responding is noteworthy for three reasons: 1) the greater perseverative responding in XXY than XY mice was observed in intact mice only, 2) this effect is conceptually similar to the relatively higher number of perseverative errors made by people with KS on the Wisconsin Card Sorting Test (Skakkebæk et al., 2013; van Rijn et al., 2012a), and 3) both XXY mice (Lewejohann et al., 2009; Lue et al., 2005) and men with KS are hypogonadal, so this result may be due to a group difference in levels of gonadal hormones.

Additionally, these differences appear to be specific increases in perseverative responding, as overall error rates (errors/trial) did not differ between genotypes. Moreover, although XY mice showed a greater across-phase increase in, and higher reversal-phase values for, *regressive* errors than XXY mice, *regressive* errors did not correlate with perseverative errors. Additionally, van Rijn et. al. (2012a) demonstrated that perseveration errors in KS men were not affected by androgen supplementation. Thus, resolving the significance for model validity of this genotype by gonadectomy interaction on perseverative errors will require further study.

Similarly, the implications of the gonadectomy-dependent learning deficit in XXY mice – but, not in XY mice – remains unclear. In van Rijn et al (2012a), perseveration errors in KS men were statistically independent of overall WCST performance and intellectual function (full-scale IQ). However, in the RLT, perseverative errors were positively correlated with trials to criterion. Moreover, the pattern of group differences in trials to criterion did not mirror those for perseverative errors. As these observations place further limitations on model validity, it appears that alternative behavioral assays of perseverative responding and learning rate, as well as additional controls for hormone levels/exposure, would be required to fully determine model validity.

### 4.3 XO vs XX mice as a model of TS

Consistent with past reports comparing 40,XO mice to XX mice on an outbred MF1 strain (Davies et al., 2005, 2007), when comparing 40,XO mice to XX mice on an inbred C57BL/6J background in the RLT, 40,XO do not show cognitive deficits. Indeed, XX mice made more premature responses than XO mice on the initially reinforced side – a result opposite of that predicted for a model of TS – and no genotype effect was observed on the side reinforced in the reversal phase. Additionally, there were no significant effects of genotype on either perseverative responding or response-holding failures (impulsivity measure), and the genotype effects on response latencies did not survive statistical correction. Lastly, although the across-phase decrease in extraneous observing responses (attention measure) was statistically significant in XX but not XO mice, within-phase genotype differences were not.

Thus, although the RLT was sufficiently sensitive to detect a number of gonadectomy effects on these measures (larger reversal cost on perseverative errors in SHAM than GDX, more premature responding in GDX than SHAM, longer response latencies in XX-GDX than XX-SHAM, and more acquisition phase extraneous observing responses in GDX than SHAM), genotype differences were either below the limit of detection or opposite of that predicted for a model of TS. However, these results do bolster the claim that the small remnant Y*^x^ chromosome in the 40,XO mice can largely rescue the cognitive deficits that may have resulted from the complete loss of a second X chromosome. That suggestion could increase the attractiveness of the steroid sulfatase gene (*STS*) as a target for TS-associated cognitive deficits (Davies et al., 2005, 2007, 2009).

A difference in copy number of *Sts* does not explain the differences between XY and XXY mice reported here as both have only one copy of *Sts* (both Y* and X^Y*^ lack *Sts*) (Burgoyne and Arnold, 2016). Nevertheless, dysregulation of *Sts* in 40, XXY mice may still occur (e.g., due to skewed X-inactivation – X vs. X^Y*^ – and/or escape from inactivation). Interest in *STS* as a contributor to phenotypic features of KS remains high because of previous studies in mice (Davies et al., 2009, 2014), and because of the association of *STS* with cognitive function and deficits in humans (Cavenagh et al., 2019; Chatterjee et al., 2016; Kent et al., 2008; Stergiakouli et al., 2011).

### 4.4 Conclusions

Greater perseveration in XXY than XY intact male mice in the RLT is a facet of inflexible responding with face validity for the greater perseverative responding in KS men than controls on the WCST. Importantly, the effects of sex hormones appeared to play a moderating role, as this genotype difference was not observed in GDX mice. These results may have implications for the use of androgens to remedy the cognitive deficits observed in KS men. Although androgen therapy does not appear to improve cognition in either KS adults (Skakkebæk et al., 2017) or children (Ross et al., 2017), little is known about the consequence of either an increased severity of hypogonadism and/or the lack of standard androgen replacement therapy on cognitive function. Our results indicate that in XXY mice, even small changes in circulating androgens can significantly change cognitive processes and learning.

