## Supplemental Figures 1-4 for "Reversal learning performance in the XY* mouse model of Klinefelter and Turner Syndromes"

### *Supplementary Figures 1-4*

#### (Reversal learning performance in the XY* mouse model of Klinefelter and Turner Syndromes)

#### SUPPLEMENTARY FIGURE 1

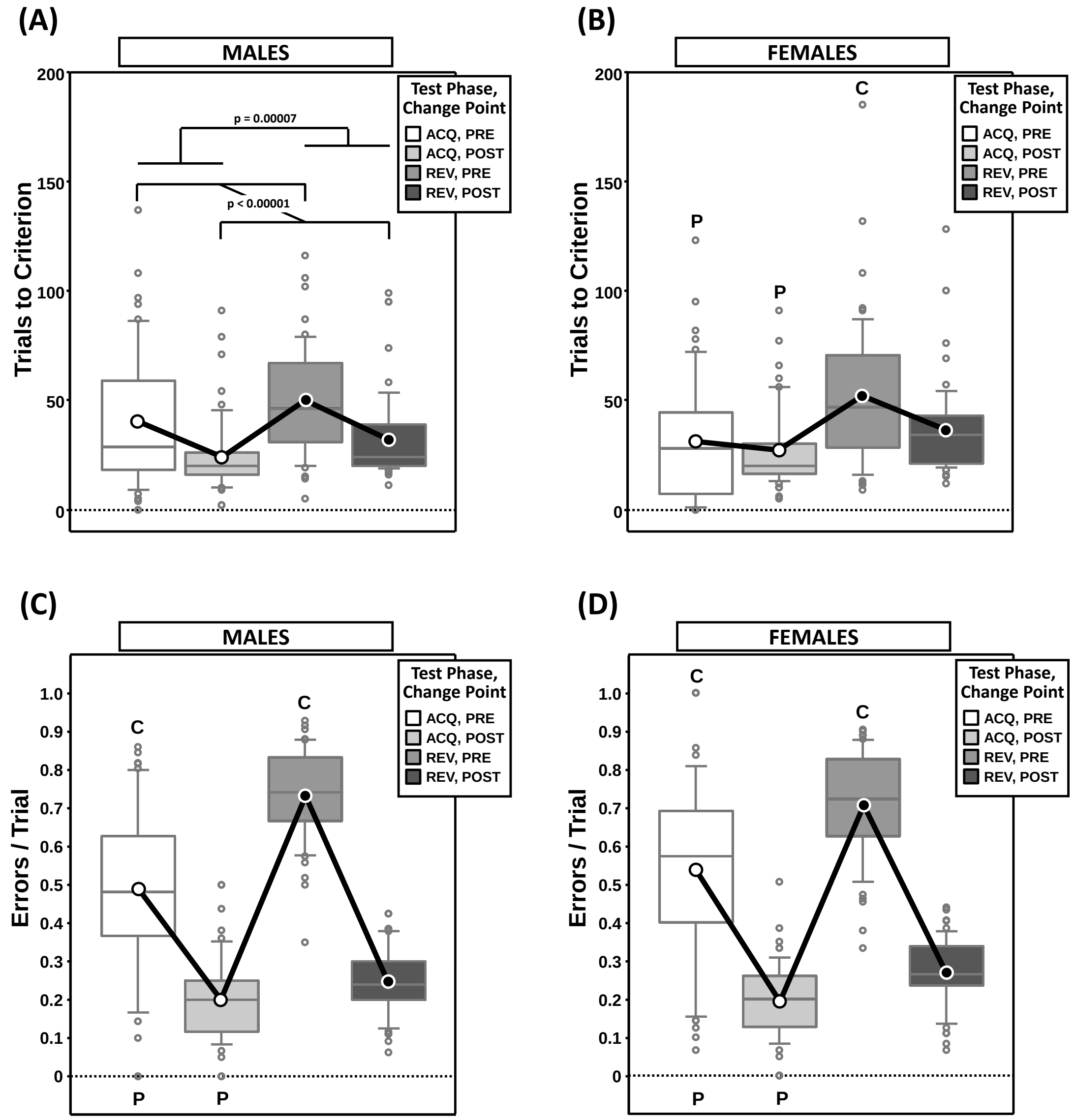


**Supplementary Figure 1:** Top panels are boxplots of trials to criterion for males **(A)** and for females **(B)** split by testing phase (Acquisition = ACQ vs Reversal = REV) and Intra-phase Change Point (PRE vs POST). For males, there was a main effect of phase ($p = 0.00007$) and change point ($p < 0.00001$), but no phase by change point interaction ($p = 0.997$). For females, there was a significant phase by change point interaction ($p = 0.041$; simple effects of phase and change point that were significant after Bonferroni correction are indicated by bold letters "P" and "C", respectively). Bottom panels are boxplots of errors per trial for males **(C)** and for females **(D)** split by Phase and Change Point. There was a significant phase by change point interaction for both males ($p < 0.0001$) and females ($p = 0.0088$). All simple effects of phase and change point were significant after Bonferroni correction (all $p < 0.001$) as indicated by bold letters "P" and "C", respectively. Boxes represent median ±quartile, whiskers extend additional $15^{th}$ of a percentile, grey open circles represent extreme values; black/white circles filled white/black represent means.

**SUPPLEMENTARY FIGURE 2**

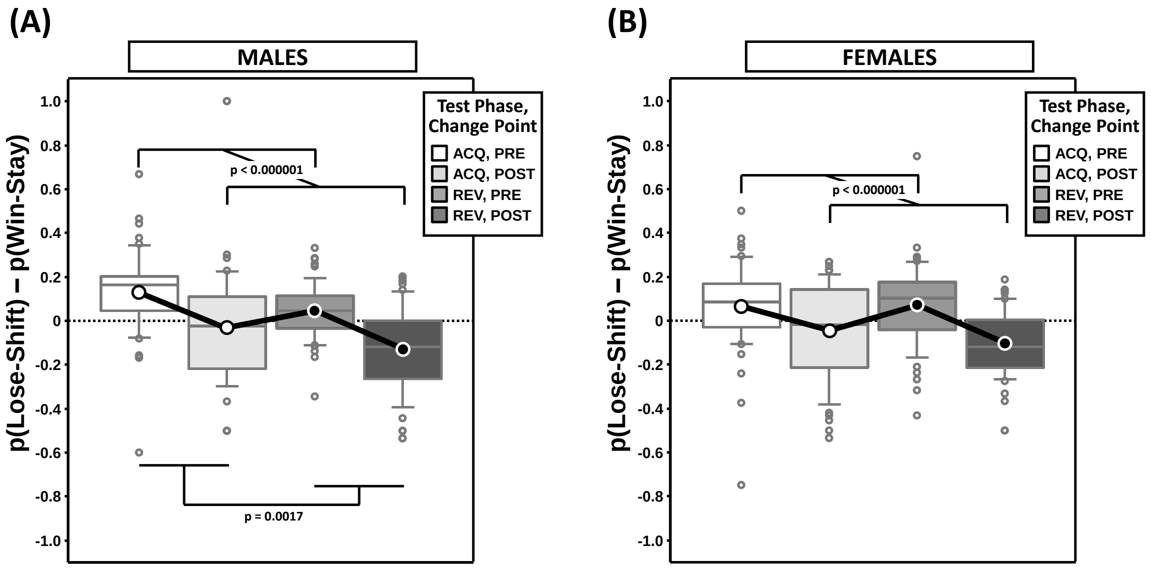


**Supplementary Figure 2:** Boxplots of the difference between the proportion of responses that were correct following either an incorrect response (**p(Lose-Shift)**; index of *flexibility*) or a correct response (**p(Win-Stay)**; index of *stability*) for males **(A)** and for females **(B)** split by testing phase (Acquisition = ACQ vs Reversal = REV) and Intra-phase Change Point (PRE vs POST). For males, there was a significant main effect of phase (p = 0.0017) and change point (p < 0.000001), but no significant phase by change point interaction (p = 0.89). For females, there was significant main effect of change point (p < 0.000001), but not of phase (p = 0.39); nor was there a significant phase by change point interaction (p = 0.16). Boxes represent median ±quartile, whiskers extend additional 15$^{th}$ of a percentile, grey open circles represent extreme values; black/white circles filled white/black represent means.

**SUPPLEMENTARY FIGURE 3**

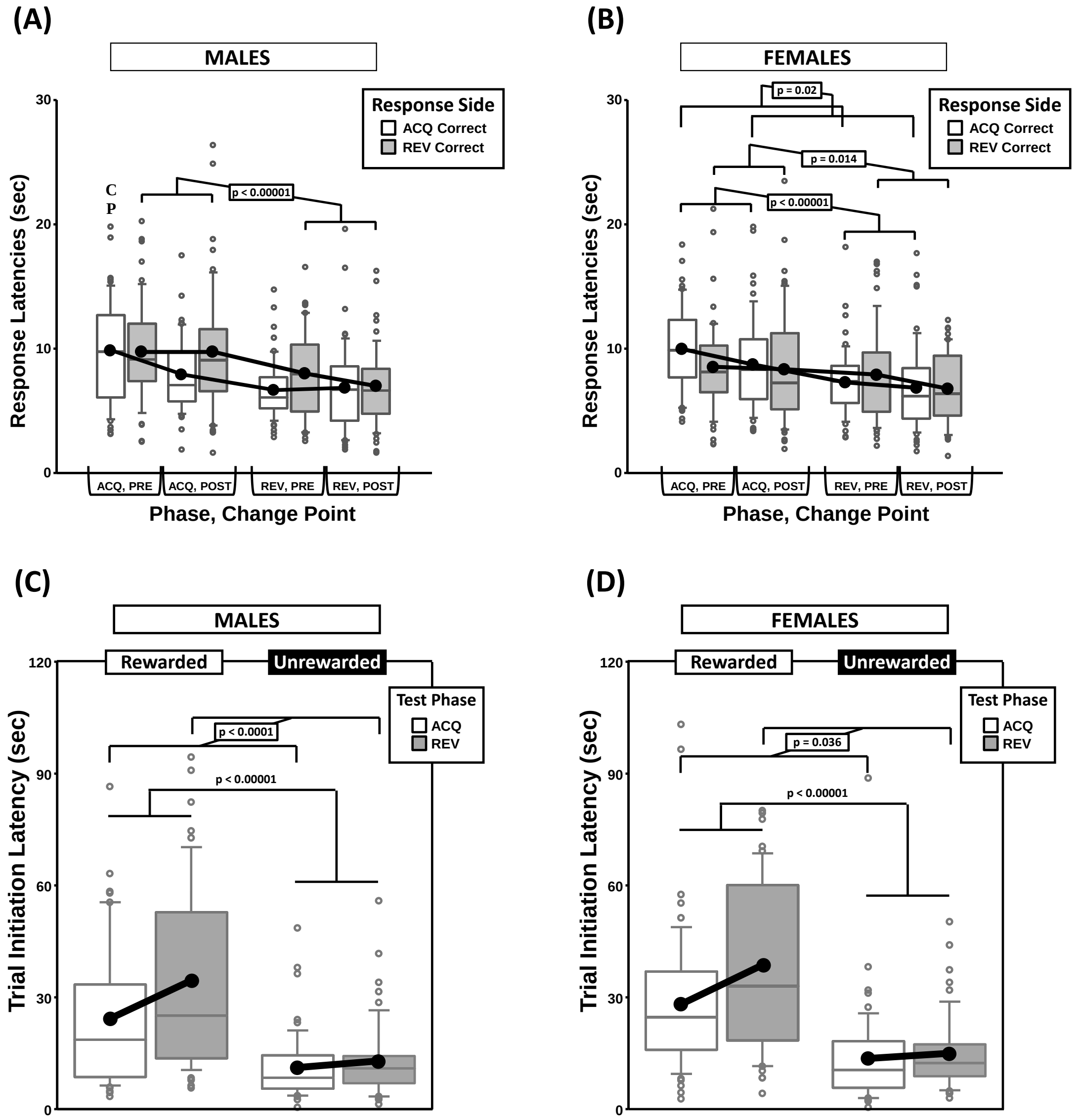


**Supplementary Figure 3:** Top panels are boxplots of response latencies to target apertures as a function of response side (correct in ACQ vs correct in REV) for males **(A)** and for females **(B)**, split by testing phase (Acquisition = ACQ vs Reversal = REV) and Intra-phase Change Point (PRE vs POST). Bottom panels are boxplots of trial-initiation latencies for males **(C)** and for females **(D)** split by testing phase and the outcome of the prior trial (rewarded vs unrewarded). Boxes represent median ±quartile, whiskers extend additional $15^{th}$ of a percentile, grey open circles represent extreme values; black circles represent means.

**SUPPLEMENTARY FIGURE 4**

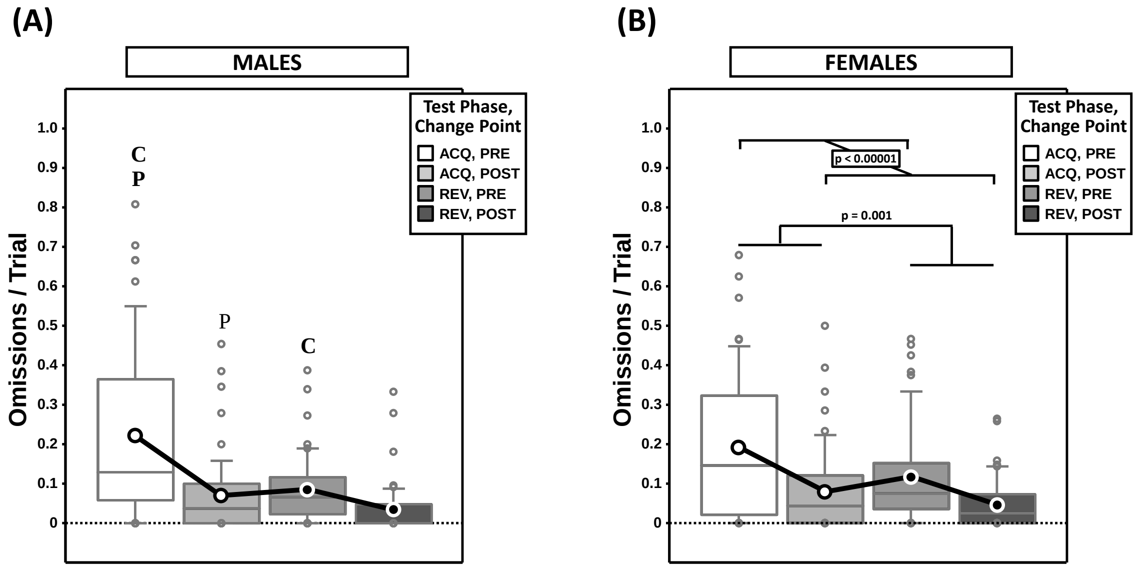


**Supplementary Figure 4:** Boxplots of omissions per trial for males **(A)** and for females **(B)**, split by testing phase (Acquisition = ACQ vs Reversal = REV) and Intra-phase Change Point (PRE vs. POST). Letters “P” and “C” indicate simple effects of Phase and Change Point ($p < 0.05$), respectively (bold letters indicate significance after Bonferroni correction). Boxes represent median ±quartile, whiskers extend additional 15$^{th}$ of a percentile, grey open circles represent extreme values; black/white circles filled white/black represent means.
